# Urban greenspace connectivity drive shifts in host assemblages and tick-borne pathogen infection

**DOI:** 10.1101/2024.11.12.623035

**Authors:** Meredith C. VanAcker, Tim R. Hofmeester, Jeffrey Zhang-Sun, Heidi K. Goethert, A. Diuk-Wasser Maria

**Author notes:** **Corresponding author:** Meredith C. VanAcker, Smithsonian’s National Zoo and Conservation Biology Institute 3001 Connecticut Ave., NW, Washington, DC, 20008, USA.

## Abstract

Habitat fragmentation is often highlighted as a driver of tick-borne disease hazard and spillover risk via reduction in biodiversity. However, habitat fragmentation can have divergent impacts on host, vector, and pathogen dynamics depending on the distribution of fragment sizes and the levels of connectivity to surrounding habitat, particularly when habitat fragments are embedded in an urban matrix. We examine how extreme habitat fragmentation influences host community composition in an urban landscape and determine its cascading impacts on *Ixodes scapularis* vector abundance and infection prevalence with human pathogenic *Borrelia burgdorferi, Babesia microti*, and *Anaplasma phagocytophilum*. We utilize camera-trapping and live mammal-trapping methods to quantify the availability of vertebrate hosts to questing larval ticks and relate relative host activity to the resulting density of nymphs and nymphal infection prevalence; the combination of these metrics determines the tick-borne disease hazard (i.e. the density of infected questing nymphs – DIN). We found that increased habitat connectivity in urban areas shifted the composition of the host community from human-adapted to forest-dependent species, species which inhabit forested habitats for all or a portion of their lifecycles. The resulting increased encounter probability between ticks and forest-dependent species increased the density of nymphs and nymphal infection prevalence with host-limited pathogens, A. *microti* and *A. phagocytophilum*, amplifying local tick-borne disease hazard. Host encounter probability of all species examined did not increase *B. burgdorferi* nymphal infection prevalence, likely due to this pathogen’s wider host range; whereas increased deer encounter probability decreased the nymphal infection prevalence of *B. burgdorferi*. These findings emphasize the importance of host identity, rather than host diversity, in shaping the heterogenous distribution of tick-borne pathogen risk in highly fragmented urban forest patches and suggest a non-linear association between disease risk and host biodiversity.

## Introduction

Tick-borne diseases are an increasing public health concern in North America, where they account for over 75% of reported vector-borne infections in the United States annually (Eisen and Paddock 2021). They continue to expand in geographic range and invade new habitats, such as urban landscapes (Rosenberg et al. 2018). Land use change (e.g. urban greening, reforestation, and habitat fragmentation) (Diuk-Wasser et al. 2021), the overabundance and range expansion of key reproductive hosts for adult ticks (Tsao et al. 2021), and climate change (Ogden et al. 2021) have driven the expansion of endemic tick species, whereas the trade of wildlife and agricultural animals has resulted in non-native tick species introductions (Rainey et al. 2018, Tsao et al. 2021). The spread of medically important tick species and pathogens infectious to humans is strongly mediated by wildlife vertebrate hosts as they provide bloodmeals for ticks, serve as pathogen reservoir hosts, and disperse feeding ticks (Tsao et al. 2021). The host community for the dominant vector of multiple tick-borne pathogens in the Eastern US, the blacklegged tick (*Ixodes scapularis*), has been well documented in the habitats that are associated with historic Lyme disease emergence: suburban and natural deciduous forests (Ostfeld et al. 2018, Mathisson et al. 2021). Now, *I. scapularis* populations are increasingly becoming established in forest remnants or newly created greenspaces in urbanized habitats where there is limited understanding of the wildlife hosts that feed, disperse, and support pathogen transmission.

Because vertebrate hosts play multiple roles in tick-borne pathogen life cycles, the structure of the host community critically determines tick-borne disease (TBD) hazard to humans, measured as the density of infected host-seeking, or questing, nymphs (DIN) because nymphs are the life stage most likely to bite and infect humans (Diuk-Wasser et al. 2021). The association between host community diversity and patterns of infectious disease (the biodiversity-disease link) has spurred significant interest and continued debate (Ostfeld and Keesing 2000, LoGiudice et al. 2008, Salkeld et al. 2013, Wood and Lafferty 2013, Civitello et al. 2015, Linske et al. 2018). A dominant theory has been the ‘dilution effect’, which proposes that the presence of additional species – in particular mesomammals that are less reservoir competent or predators of highly competent rodents, in a community directly or indirectly inhibits parasite transmission, resulting in a negative relationship between species diversity and disease (Keesing et al. 2006). While the diversity of roles played by different hosts is widely recognized and previous work has demonstrated that vertebrate host species vary in their ability to host ticks and acquire and transmit tick-borne pathogens (Donahue et al. 1987, Telford III et al. 1988, LoGiudice et al. 2003, Vuong et al. 2014), the linkages between landscape structure, host community composition, and pathogen amplification have been strongly debated and have not been empirically assessed.

Although changes in the host community are considered the direct driver of changes in the TBD hazard, anthropogenic habitat fragmentation has often been used as a proxy for biodiversity under the critical assumption that fragmentation leads to host community enrichment of highly reservoir competent small mammalian hosts, as seen in agricultural landscapes in the Midwest U.S. (Nupp and Swihart 1998, 2000). Studies using forest fragmentation as a proxy for TBD hazard have found conflicting results (Allan et al. 2003, Brownstein et al. 2005, LoGiudice et al. 2008, Diuk-Wasser et al. 2012, Zolnik et al. 2015) where the strongest impact on TBD hazard is claimed to occur in very small forest patches (∼1 ha) (Allan et al. 2003, Diuk-Wasser et al. 2021), though another study found that too few ticks were collected in small patches (2.7+- 3.1 ha, (mean +- SD)) to estimate pathogen prevalence (LoGiudice et al. 2008). This suggests that tick establishment thresholds could limit potential impacts of fragmentation on the host community composition (LoGiudice et al. 2008). Because habitat patches of this small size do not capture the home ranges of most TBD hosts (Randolph and Dobson 2012, VanAcker et al. 2023), there’s a need to understand the role of the non-forest habitat matrix in structuring host communities at a landscape scale (Diuk-Wasser et al. 2021). The only study assessing the role of landscape structure on TBD hazard conducted in an urban area in the U.S. found that, in fact, connectivity across green spaces was positively associated with TBD hazard (VanAcker et al. 2019), consistent with studies in Europe that link urban green space connectivity with tick presence and increased tick densities (Heylen et al. 2019, Hansford et al. 2023).

The ecological contexts within which studies assess the effects of fragmentation on host community assembly are critical to consider when drawing comparisons between findings. Previous studies utilized patch scale findings to infer the effect of fragmentation on landscape scale processes (Allan et al. 2003, Brownstein et al. 2005, Diuk-Wasser et al. 2012, Zolnik et al. 2015). While other studies utilized host assembly results from forest patches set within an agricultural matrix (Nupp and Swihart 1998, Krohne and Hoch 1999, Rosenblatt et al. 1999) to infer host community assembly in forest patches within a matrix of suburban development (Allan et al. 2003, Keesing et al. 2006). However, agricultural and urban landscapes offer fundamentally different resources and levels of matrix permeability for wildlife hosts to move between patches making the ecological contexts challenging to compare. Additionally, while most studies assess fragmentation considering the size of the habitat patch itself, the total amount of habitat surrounding the sampling sites may be the key driver of species richness (Fahrig 2013). To address these limitations, we used a landscape-based approach to assess mammalian host communities in forested habitat patches surrounded by an urban matrix allowing us to examine how increasing habitat amount and levels of connectivity impact host community assembly at a landscape scale.

*I scapularis* is a three-host generalist feeder and can acquire and transmit zoonotic pathogens among a wide suite of mammals and birds during the tick’s bloodmeal (Eisen and Eisen 2018, Halsey et al. 2018). Immature stages feed on a wide range of mammalian and avian hosts, including white-tailed deer (Huang et al. 2019), and adult *I. scapularis* feed almost exclusively on white-tailed deer (*Odocoileus virginianus,* hereafter deer). In contrast to their generalist vector, *Ixodes scapularis*-borne pathogens vary in the range of hosts that serve as competent reservoirs. The most widespread tick-borne pathogen in the U.S. is *Borrelia burgdorferi* sensu stricto, the causative agent of Lyme disease, which is transmitted horizontally between *I. scapularis* and reservoir hosts that vary in their level of competence for *B. burgdorferi* and their ability to provide bloodmeals to immature ticks (LoGiudice et al. 2003, Brisson and Dykhuizen 2004, Brunner et al. 2008). *Ixodes scapularis* also vectors agents of babesiosis and anaplasmosis, which are increasing in human incidence (Rosenberg et al. 2018). Babesiosis in the Eastern U.S. is caused by infection with a protozoan parasite, *Babesia microti.* White footed mice, *Peromyscus leucopus*, are considered the primary reservoir host for *B. microti* (Spielman et al. 1981, Mather et al. 1990, Goethert et al. 2018), although other mammals can serve as reservoir hosts (Hersh et al. 2012). *B. microti* can also be vertically transmitted in *P. leucopus* (Tufts and Diuk-Wasser 2018, 2021), strengthening the role of *P. leucopus* in maintaining the *B. microti* transmission cycle, particularly in isolated habitat patches. The etiologic agent of anaplasmosis in the United States is a rickettsial bacterium, *Anaplasma phagocytophilum*, of which there are two major variants that have been associated with different reservoir hosts; *P. leucopus* is the main reservoir host for the Ap-ha variant (Massung et al. 2002) and deer are the primary reservoir host for the Ap-variant 1, which lacks the ability to infect *P. leucopus* and has not been linked to human infection (Massung et al. 2003b).

In this study, we quantified host availability for questing *I. scapularis* in a multi-host community to determine how the density and infection of ticks varies with host species activity. We use host availability to ticks as a metric measured through passage rate estimates from camera trapping meso-mammals and abundance estimates from live trapping small mammals. We assessed how the area and connectivity of habitats differentially affected the relative abundance of host species (‘host identity hypothesis’, MacDonald et al. 2022) as well as host diversity (‘dilution effect’) for questing ticks and the resulting pathogen prevalence in local tick populations. During 2018 and 2019, we employed camera trapping, live trapping, and tick sampling in natural habitats within public parks in a borough of New York City (NYC), Staten Island, coextensive with Richmond County, NY, to address the following questions: (1) How do area and connectivity of green spaces in an urban matrix affect host availability for *Ixodes scapularis* ticks? (2) Does the activity of host species relate to nymphal tick abundance? and (3) How does the probability of a larva encountering a host of a particular species, relative to others in the community, relate to the prevalence of pathogens in questing nymphs?

## Methods

### Sampling design: tick collection and small mammal trapping

Each year, we selected eight forest patches (> 6 ha in area) that spanned Staten Island, a borough in NYC, and varied in their level of forest connectivity and park area (minimum area = 6.85 ha and maximum area = 307 ha) (VanAcker et al. 2019). At each site we established a sampling grid organized as a 10 x 5 rectangular array with 10 meters between the nodes (*n* =50), equivalent to ten 40 m transects. Five sites were sampled both years, three sites were sampled in 2018 only, and three sites were sampled in 2019 only, totaling 11 unique sites. The density of *I. scapularis* nymphs was estimated by dragging a 1 m^2^ cloth along the established transects, stopping every 10 m at the grid nodes to avoid tick fall-off and collecting all ticks in a unique tube per transect (Daniels and Fish 1990). Sampling was conducted every 2 weeks (referred to as sessions) between late-May and the end of August/early September (2018: June 3 – August 23; 2019: May 26 – September 10) in 2018 and 2019. Adult and nymphal ticks were collected in 90% ethanol or collected live to be frozen at -80° C. Larval ticks were collected using clear packaging tape. All ticks’ species, life stage, and counts were confirmed in the lab using taxonomic keys (Keirans and Clifford 1978, Durden and Keirans 1996).

Ticks were kept in ethanol or in the freezer until DNA was extracted using the DNeasy Blood and Tissue Kit (Qiagen, Valencia, CA, USA) according to manufacturer’s recommendations (DNA extraction details in Appendix S1). All DNA extractions were shipped on dry ice to the Cummings School of Veterinary Medicine at Tufts University where the samples were screened by H. Goethert according to published methods (Goethert et al. 2021). Up to ∼200 *I. scapularis* ticks collected from each site were screened individually for infection with *B. burgdorferi*, *B. microti*, and *A. phagocytophilum* using a multiplex real-time PCR assay with primers and probes that were previously published (Tokarz et al. 2017) (PCR assay details in Appendix S1: Table S1(a)).

Mark-recapture of small mammal hosts via live trapping was conducted over two consecutive days during each session. Sherman traps were placed at each of the grid’s nodes at 10 m intervals (*n* = 50 traps/grid). Upon capture, all small mammals were ear-tagged, aged, sexed, weighed, and screened for ticks before release at the point of capture. One 1 mm ear punch was taken from each mouse and stored in 100% EtOH until returning to the lab and a blood sample was taken and dried on Whatman FTA cards (Fisher Scientific, Pittsburg, PA, USA). Blood samples were prepped for DNA extraction using the Qiagen DNeasy Blood and Tissue kit following the manufacturer’s protocol (Qiagen, Valencia, CA, USA) and DNA was extracted using the QIAamp DNA extraction robot (DNA extraction details in Appendix S1). The DNA extracts from blood were screened for *B. microti* using qPCR specific primers and probes (Appendix S1: Table S1(b)). We used the mark-recapture data for *P. leucopus* to estimate mouse relative abundance, referred to as minimum number alive (MNA), at each site according to Krebs methodology (Krebs 1966), shown to provide reliable estimates of abundance with capture heterogeneity (Davis et al. 2003). To estimate MNA, the data was divided into the number of mice caught during each session (time *t*) and the number of individuals not caught during that session but known to be present due to subsequent captures. Then, the individuals caught prior and following time *t* were summed to estimate the MNA value. This index was used to model the relationship between *I. scapularis* density and infection and mouse abundance.

### Camera deployment

We deployed two Moultrie M-50 game cameras at each of the eight sites so that 16 cameras were active at any given time. The cameras were rotated every 2 weeks to match the tick collection and live trapping schedule and were active throughout the duration of tick and small mammal sampling. We divided each grid into two square sub-plots and deployed one camera within each sub-plot. We determined camera locations by generating random coordinates within the perimeter of each sub-plot with a distance of at least 30 m between the two coordinates to prevent sampling overlap (Hofmeester et al. 2017c). The cameras were mounted to a tree closest to the random coordinate 20-40 cm off the ground to mimic the height of questing ticks (Tietjen et al. 2020) and following the methods of Hofmeester et al. 2017 (additional camera setting details in Appendix S1). The detection distance was estimated by having researchers walk away from the camera’s point of view until the camera no longer detected the researcher and the distance (m) was recorded. Detection distance was estimated each time the camera was relocated. We placed flags within the camera’s point of view every 2.5 m to establish a minimum of five distance classes summing to 12.5 m total, the minimum distance shown to reduce bias due to body mass and vegetation in estimating effective detection distance (Hofmeester et al. 2017c). Theft and camera malfunction led to some loss of data from three cameras, however because of the study design, data was captured from at least one camera at each site for all timepoints throughout the study duration resulting in continuous representation of the grids. There were five sites that were sampled across both years, the camera locations across years were within the same grid area but different random GPS locations were used across the study period.

### Photo screening

Camera trap photos from 2018 and 2019 were sorted and screened using the software *Camelot* (v1.5.6 and v1.6.3) by the same observer within year but different observers between years. We maintained consistent qualifiers for species identification, life stage, and animal sex (deer only). In any case of uncertainty due to image quality or an obstructed view, we declared an unknown sex and/or unknown life stage. If species could not be identified in photos, photos were not included in analysis.

We used a 30-minute interval for the threshold of independence between individuals of the same species in photos. Thus, any other photo taken within 30 minutes of the first photo that captured an individual of the same species was counted as one observation. However, if the photographed animals could be distinguished by differences in sex or life-stage (only valid for deer), we would mark the individuals as independent, and this would qualify as two observations. We assigned each photo to a distance interval by observing which distance flag the animal was closest to when crossing the midline of the camera’s line of sight.

### Estimating host passage rates and encounter probabilities

We drew on previously published methods (Rowcliffe et al. 2011, Hofmeester et al. 2017c, 2017b) to estimate the passage rates and encounter probability of each host species across all sites where camera trapping took place during 2018 and 2019. The method of using camera traps as proxies for animal detection by ticks relies on the idea that animal detection by cameras is a function of the frequency and proximity of passage – similar to that of host detection by ticks (Hofmeester et al. 2017b).

First, we estimated effective detection distance (EDD) – the distance where the count of animals detected further away is estimated to be equivalent to the count of animals missed closer – for each host species that had > 20 independent photos over each season. To do this we parameterized the following equation:

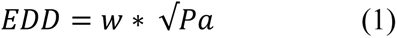

where *w* is the truncation distance and *Pa* is the expected probability of detection within distance *w* from the camera’s point of view. We used the farthest distance the animal species was detected as the truncation distance (12.5 m or farther) and estimated *Pa* using mark-recapture distance sampling detection models in the *mrds* (Laake et al. 2021) package in R. To estimate *Pa*, the distribution of passages across the distance intervals was used to fit a detection probability function following point model techniques (Buckland et al. 2001). A point model assumes the distribution of the number of detections is similar to the probability of detection multiplied by distance to correct for the increase in detection area with distance and has been shown to describe detection by sensors well (Rowcliffe et al. 2011, Hofmeester et al. 2017c). We compared the fit of two detection probability models, a hazard rate and a half-normal model, without covariates on single species data. We used AIC comparison and retained the model with the lowest AIC as the best fit to the data (Buckland et al. 2015) and estimated effective detection distance using equation 1 where the *Pa* parameter was informed by model outputs.

To determine hosts’ activity by camera traps, referred to as a species’ passage rate, *Pi*, we used an equation where animal passage rate is proportional to the rate of contact between animals and ticks as animals are assumed to walk randomly past questing ticks:

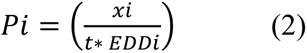

*x_i_* is the number of independent passes of species *i*, *t* is the number of days each camera was active, and *EDD_i_* is the effective detection distance of species *i* (in meters) (Hofmeester et al. 2017b). Passage rate m^-1^ d^-1^ can be interpreted as an index of local species activity defined by the number of times an animal passes in front of the camera per day standardized by detectability differences, such as differences due to body mass (Hofmeester et al. 2017a). Because we were also interested in a tick’s probability of encountering a particular host species *relative* to the community of hosts at each site for pathogen transmission, we estimated the encounter probability (EP) of each host species at any given site (Takumi et al. 2019):

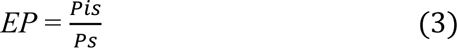

*Pis* indicates the passage rate of host species *i* at site *s* and *Ps* is the sum of the passage rate of all host species at site *s*.

### Host passage models

All statistical modeling was performed in R (version 4.0.1) (Team 2022). We estimated mammalian host diversity at each site using the Shannon index (Shannon 1948) where the number of independent observations of a particular species/night was used as *n_i_*. We used data from the NYC Green Spaces map (NYC 2015) to estimate the total area in ha of each park where the grids were located. To estimate a park’s level of connectivity, we estimated each park’s mean and median cumulative current, the total current flowing through the landscape, using the Omniscape program (Landau et al. 2021) (layer construction details in Appendix S1). We ran Omniscape with four different moving window radii ranging from 120 m to 1830 m and assessed the fit of the mean and median cumulative current at each sliding window size by running generalized linear mixed models (GLMMs) with the negative binomial family to predict the density of nymphs and GLMMs with the poisson family to predict deer passage rate; both models used year as a random effect. By estimating connectivity using a moving window, we explicitly examine the ‘habitat amount’ hypothesis (Fahrig 2013) where the amount of habitat within the moving window radii is integrated into the site’s connectivity estimate. We assessed the relationships between park area, connectivity, and Shannon index using gaussian generalized linear models (GLMs). We examined the drivers of variation in host species activity using GLMMs for deer, raccoon, and squirrels – the most abundant mammalian host species detected. We included the total area of the park, Shannon index, and each park’s connectivity value (120 m sliding window) as fixed effects while year was used as a random effect. We estimated the total number of cumulative independent passes for each species at each site per year (*x_i_/EDD_i_* from eqn. 2) as the response variable in the models. We used the R package *glmmTMB* (Brooks et al. 2017) with a Poisson distribution and included an offset for the log number of total days cameras were active (*t* from eqn. 2). Additionally, we used negative binomial GLMs to examine the relationship between mouse abundance and park area, connectivity, and Shannon index while utilizing year as a random effect. Finally, we assessed the relationships between park area, connectivity, and Shannon index using gaussian GLMs.

### Host community structure

We characterized host community structure by examining the correlation between host species’ activity and their putative driving factors (park connectivity, park area, and Shannon index) by running a standardized Principal Component Analysis (PCA) using the *factorextra* (Kassambara and Mundt 2020) package in R. PCA reduces the dimensionality of a data set by minimizing the residual variance across data to create a visualization of summary axes called principal components. We ran separate PCAs for 2018 and 2019, each including all host species that had > 20 independent observations. We plotted the top two principal components that accounted for the highest variance explained and overlaid the parks’ connectivity, area, and Shannon index to examine how each dimension correlated with the site characteristics.

### Tick abundance and infection prevalence models

We examined the associations between host activity and park characteristics with the density of nymphs (DON), the relationship between park characteristics and DIN, and host encounter probabilities with the infection prevalence of nymphs (NIP) for all three pathogens. We estimated DON by summing the counts of *I. scapularis* nymphs per transect and dividing by the tick dragging transect length completed during the maximum nymphal activity season (sessions 1 – 5 for both 2018 and 2019: June 3 – August 9, 2018, and May 26 – August 1, 2019). We modeled the relationship between DON and park characteristics using negative binomial GLMs with a “logit” link.

We ran Spearman’s correlation on the nymph counts/transect and host activity variables for each year separately prior to modeling and only modeled covariates together that had a correlation value less than 0.70. All covariates used in these multivariate models were standardized using the “scale” function. We used negative binomial GLMs with a “logit” link to assess the relationship between host activity and DON. We used the “dredge” function from the *MuMIn* (Bartoń 2009) package to examine all possible subsets of the global models and reported the models within 2 ΔAIC from the lowest AIC model. We estimated the variance inflation factor (VIF) of each model to ensure all variables had a VIF < 4 (Fox 2015). We modeled the association between DON with host activity using a one-year lag because the life of *I. scapularis* spans over two years and successful host feeding as a larva (2018) relates to abundance the following year (2019) in the nymphal stage.

At sites where > 40 nymphs were collected and screened for pathogens, we estimated NIP by dividing the number of infected nymphs out of the total number of screened nymphs separately for each pathogen and site. Due to sample size limitations, we were not able to use a lagged year analysis to predict NIP, thus data from the two years was combined and a random effect for year was included. We used a binomial GLM to model the relationship between host encounter probabilities and NIP for *B. burgdorferi, B. microti*, and *A. phagocytophilum* using deer and raccoon encounter probabilities and mouse abundance. High collinearity between squirrel and raccoons prevented the inclusion of squirrel encounter probability in the models. To estimate DIN, we multiplied DON and NIP from each site and assessed the relationship between DIN and park characteristics using negative binomial GLMs with a “logit” link. Finally, we examined the occurrence of pathogen coinfection within nymphal ticks and assessed whether coinfection occurred more than expected based on the pathogens’ individual prevalence in ticks using a Chi-squared test.

## Results

### Tick collection and pathogen infection and coinfection

The mean density of nymphs in 2018 was 59/1000 m (SD ±61.96) with a minimum of 0 and a maximum of 171/1000 m. In 2019, the densities declined to a mean nymphal density of 25/1000 m (SD ±33.79) with a minimum of 1/1000 m and a maximum of 102/1000 m.

We extracted DNA from a total of 1056 ticks in 2018 and 480 in 2019 and screened for *B. burgdorferi*, *B. microti*, and *A. phagocytophilum* (Appendix S1: Table S2). Ticks from sites where there was a minimum of 40 *I. scapularis* nymphs collected per year were screened for the presence of *B. burgdorferi*, *B. microti*, and *A. phagocytophilum* resulting in 10 out of 18 total site-years (Table 1.). Average NIP across sites was 21% for *B. burgdorferi* in 2018 and 22% in 2019, *B. microti* NIP was 10% in 2018 and 13% in 2019, and *A. phagocytophilum* NIP was 6% in 2018 and 18% in 2019 (Table 1.). The proportion of nymphs coinfected with *B. burgdorferi* and *B. microti* was 5% in 2018 and 2019. The proportion of nymphs coinfected with either *B. burgdorferi* and *A. phagocytophilum* or *B. microti* and *A. phagocytophilum* was <1% in 2018 and 1% in 2019. The increase in *A. phagocytophilum* NIP between 2018 and 2019 was higher than expected by chance based on a Chi-squared test (*X*^2^ = 38.77; P < 0.0001) while the increase in *B. burgdorferi* and *B. microti* NIP was not significant. We found the coinfection rate with *B. burgdorferi* and *B. microti* was higher than expected by chance, while the other observed coinfection rates were not different than expected by chance (all infection results in Appendix S1: Table S2.).

**Table 1.**
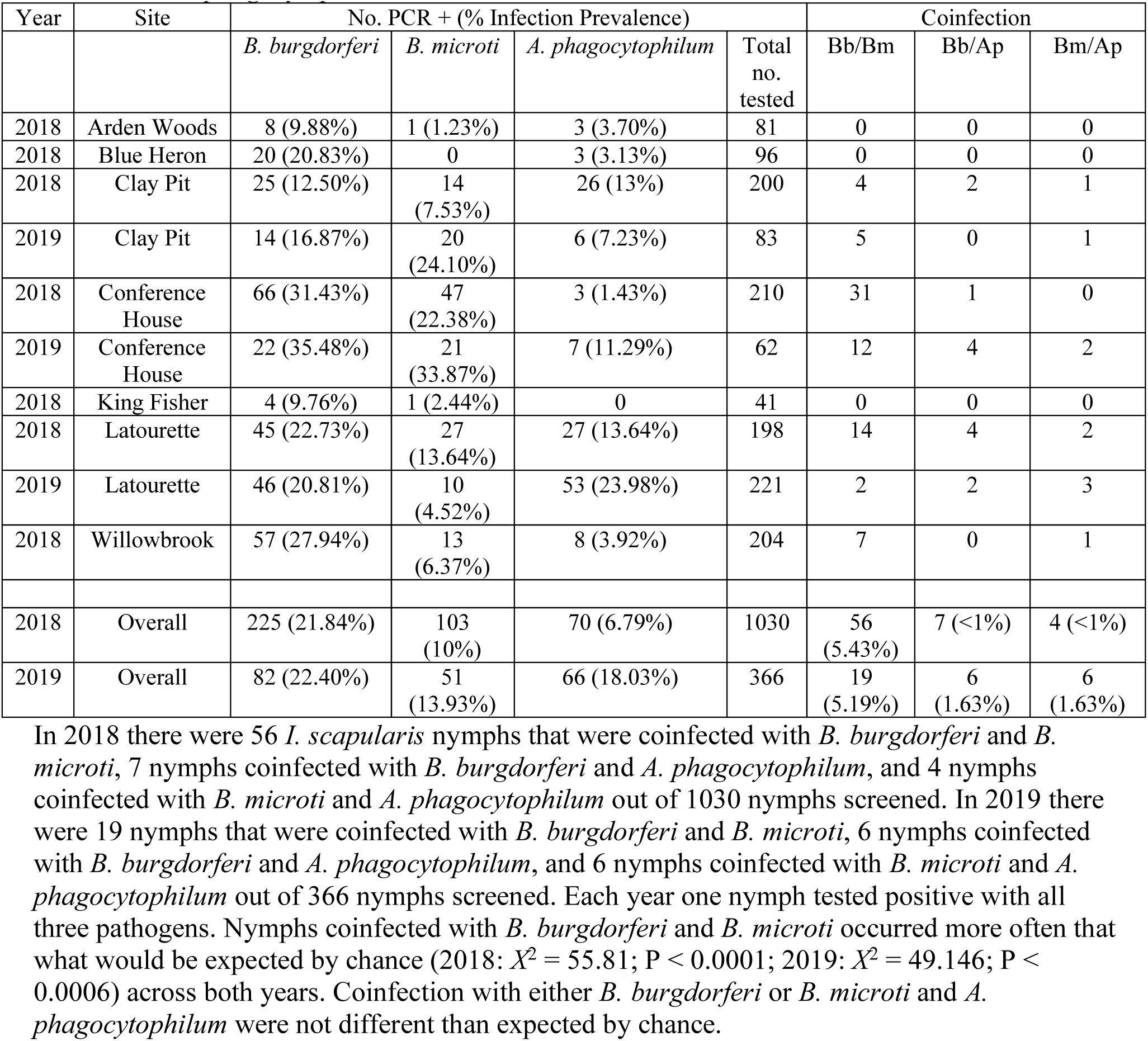
*I. scapularis* nymphal infection prevalence and coinfection for *B. burgdorferi*, *B. microti*, and *A. phagocytophilum* in 2018 and 2019.

| Year | Site | No. PCR + (% Infection Prevalence) |  |  |  | Coinfection |  |  |
| --- | --- | --- | --- | --- | --- | --- | --- | --- |
|  |  | *B. burgdorferi* | *B. microti* | *A. phagocytophilum* | Total no. tested | Bb/Bm | Bb/Ap | Bm/Ap |
| 2018 | Arden Woods | 8 (9.88%) | 1 (1.23%) | 3 (3.70%) | 81 | 0 | 0 | 0 |
| 2018 | Blue Heron | 20 (20.83%) | 0 | 3 (3.13%) | 96 | 0 | 0 | 0 |
| 2018 | Clay Pit | 25 (12.50%) | 14 (7.53%) | 26 (13%) | 200 | 4 | 2 | 1 |
| 2019 | Clay Pit | 14 (16.87%) | 20 (24.10%) | 6 (7.23%) | 83 | 5 | 0 | 1 |
| 2018 | Conference House | 66 (31.43%) | 47 (22.38%) | 3 (1.43%) | 210 | 31 | 1 | 0 |
| 2019 | Conference House | 22 (35.48%) | 21 (33.87%) | 7 (11.29%) | 62 | 12 | 4 | 2 |
| 2018 | King Fisher | 4 (9.76%) | 1 (2.44%) | 0 | 41 | 0 | 0 | 0 |
| 2018 | Latourette | 45 (22.73%) | 27 (13.64%) | 27 (13.64%) | 198 | 14 | 4 | 2 |
| 2019 | Latourette | 46 (20.81%) | 10 (4.52%) | 53 (23.98%) | 221 | 2 | 2 | 3 |
| 2018 | Willowbrook | 57 (27.94%) | 13 (6.37%) | 8 (3.92%) | 204 | 7 | 0 | 1 |
| 2018 | Overall | 225 (21.84%) | 103 (10%) | 70 (6.79%) | 1030 | 56 (5.43%) | 7 (<1%) | 4 (<1%) |
| 2019 | Overall | 82 (22.40%) | 51 (13.93%) | 66 (18.03%) | 366 | 19 (5.19%) | 6 (1.63%) | 6 (1.63%) |

In 2018 there were 56 *I. scapularis* nymphs that were coinfecting with *B. burgdorferi* and *B. microti*, 7 nymphs coinfecting with *B. burgdorferi* and *A. phagocytophilum*, and 4 nymphs coinfecting with *B. microti* and *A. phagocytophilum* out of 1030 nymphs screened. In 2019 there were 19 nymphs that were coinfecting with *B. burgdorferi* and *B. microti*, 6 nymphs coinfecting with *B. burgdorferi* and *A. phagocytophilum*, and 6 nymphs coinfecting with *B. microti* and *A. phagocytophilum* out of 366 nymphs screened. Each year one nymph tested positive with all three pathogens. Nymphs coinfecting with *B. burgdorferi* and *B. microti* occurred more often than what would be expected by chance (2018: $X^2 = 55.81$ ; $P < 0.0001$ ; 2019: $X^2 = 49.146$ ; $P < 0.0006$ ) across both years. Coinfection with either *B. burgdorferi* or *B. microti* and *A. phagocytophilum* were not different than expected by chance.

### Mesomammal camera and live trapping

In 2018, we ran a total of 1081 camera nights and, in 2019, we ran a total of 1582 camera nights distributed across 16 cameras at 8 sites, five sites of which were monitored during both years. Cameras detected 27 avian and mammalian species in 2018 and 45 avian and mammalian species in 2019. We focused on six mammalian species documented to host *I. scapularis* (domestic cat – *Felis catus*; Eastern gray squirrel – *Sciurus carolinensis*; groundhog – *Marmota monax*; raccoon – *Procyon lotor*; Virginia opossum – *Didelphis virginiana*; white-tailed deer – *O. virginianus*). We did not include bird species in our analyses as prior studies have shown bird detection reliability declines greatly following a 60 cm camera-subject distance (Ortmann and Johnson 2021). Deer, raccoon, and squirrel were found across all 11 unique sites and deer showed the highest activity. Due to low activity (opossum and groundhog) or little evidence for the species to be an important host to *I. scapularis* (cats; (Tufts et al. 2021)), opossums, groundhogs, and cats were not included in the models, although passage rates were estimated for each species (Appendix S1: Figure S1). The hazard rate function was the best fit to all species detection data, except for deer in 2019 where the half-normal detection function was a better fit indicated by AIC comparison (Table 2). The EDD ranged from 8.32 to 9.69 m in 2018 and 5.72 to 7.92 m in 2019 across cat, deer, opossum, raccoon, and squirrel (Table 2 and Appendix S1: Figure S2). We captured a total of 86 unique *P. leucopus* individuals in 2018 and 144 in 2019. The MNA estimates ranged from 2 – 39 in 2018 and from 4 – 90 individuals in 2019 (Appendix S1: Figure S3).

**Table 2.** Model fits as indicated by Akaike Information Criterion (AIC) scores from detection function analyses using the half-normal and the hazard rate functions by host species and year.

| Species | Year | Half Normal AIC | Hazard Rate AIC | EDD (m) |
| --- | --- | --- | --- | --- |
| Raccoon | 2018 | 848.37 | **797.87** | 9.69 |
| Raccoon | 2019 | 1876.32 | **1839.41** | 7.20 |
| Opossum | 2018 | 97.21 | **90.97** | 8.87 |
| Opossum | 2019 | 506.58 | **463.04** | 6.50 |
| Squirrel | 2018 | 1897.56 | **1712.82** | 8.32 |
| Squirrel | 2019 | 2498.56 | **2341.03** | 7.80 |
| Domestic cat | 2018 | 79.76 | **78.11** | 8.61 |
| Domestic cat | 2019 | 834.81 | **810.62** | 5.72 |
| White-tailed deer | 2018 | 5456.13 | **5405.05** | 9.60 |
| White-tailed deer | 2019 | **18763.63** | 19026.83 | 7.92 |
| Groundhog | 2019 | 162.63 | **161.42** | 6.66 |

The estimated effective detection distances (EDD) are listed for each host species and year.

### Host activity models

The scale sensitivity analysis of Omniscape’s moving window radius resulted in the median cumulative current flow within 120 m radii as the best scale to predict deer activity and the mean cumulative current flow within 1830 m radii as the best scale to predict DON (Appendix S1: Figure S4). We did not find a significant relationship between park area and connectivity (120 m) or Shannon diversity index but did observe a significant negative relationship between Shannon diversity index and park connectivity (ß = -0.12, p = 0.005; Appendix 1: Figure S5). We used connectivity within 120 m window to model all host related processes and connectivity within 1830 m window to model all tick and pathogen dynamics. Deer activity was higher in more connected (ß = 0.12, *p* < 0.01), smaller (ß = -0.29, *p* < 0.001), and less host diverse (ß = - 0.26, *p* < 0.001) parks; raccoon activity was higher in smaller (ß = -0.37, *p* = 0.02), and more host diverse (ß = 0.52, *p* < 0.001) parks and park connectivity was not present in the best fit raccoon model. Squirrel activity was higher in less connected (ß = -0.47, *p* < 0.001) and larger parks (ß = 0.18, *p* = 0.03), and host diversity within parks was not present in the best fit model (Figure 1, Appendix S1: Table S3). Mouse abundance (white-footed mouse MNA) was higher in larger (ß = 0.49, *p* = 0.002) and more isolated parks (ß = -0.70, *p* < 0.001) and Shannon diversity index was not present in the model with the lowest AIC (Appendix S1: Table S4).

**Figure 1.**
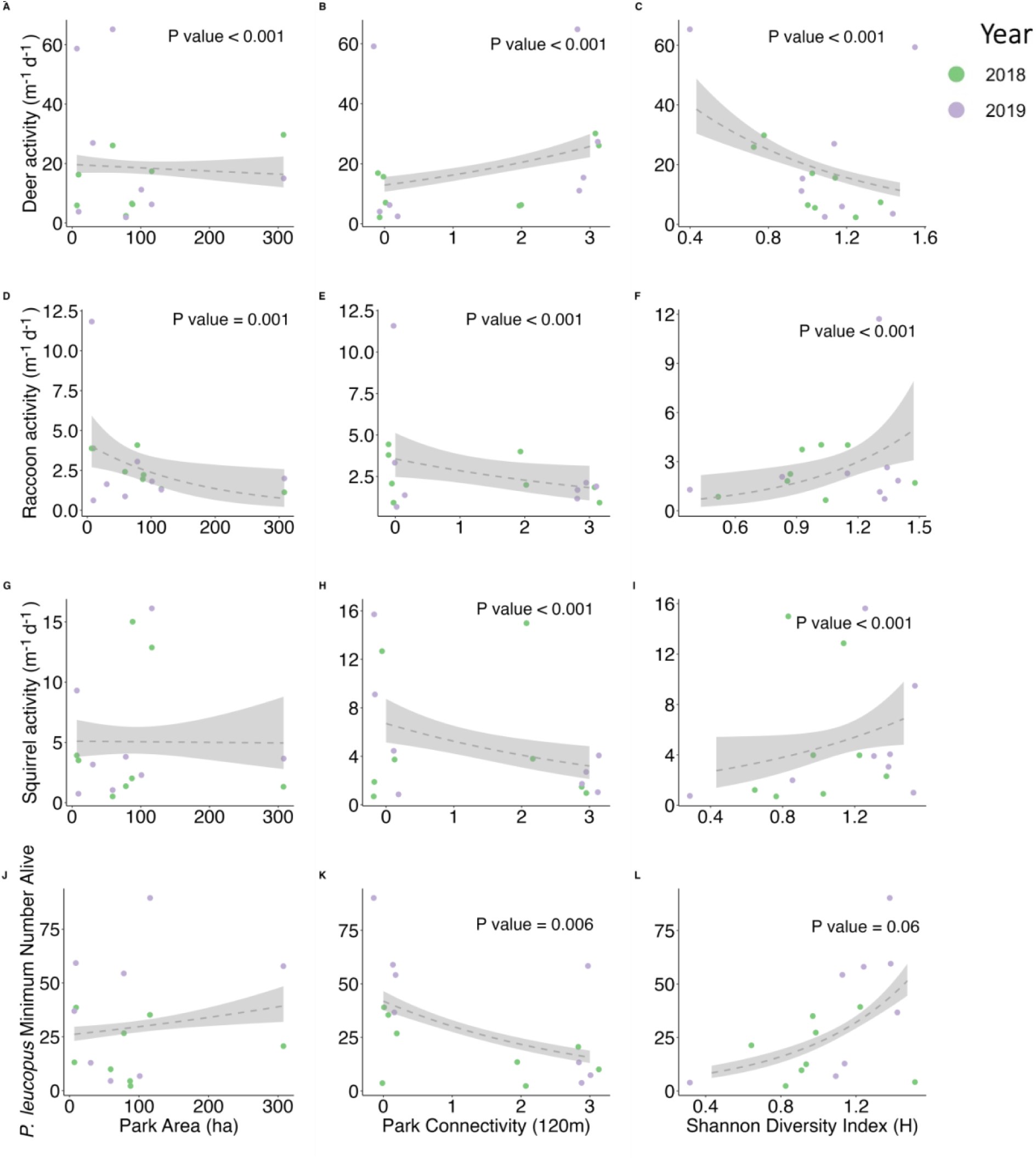
Univariate relationships between deer, raccoon, and squirrel activity and the minimum number alive for *P. leucopus* with park area (ha), connectivity, and Shannon’s index. The points show site species host activity, and the color indicates the year. Shannon’s index is the only predictor variable that varied by year. Dashed lines show the model fit of each univariate Poisson model using the GLM smoothing method and the gray bands indicate 95% confidence intervals. The P-values are included for the significant or near significant models. Activity is indicated by passage rate whose units are in m^-1^ d^-1^ and data is presented as an unstandardized estimate.

### Host community structure

The first two axes of the 2018 PCA explained 69% of the variance; opossum, raccoon, and cat passage rates contributed most significantly to principal component (PC) 1 summing to 72%, while deer, mice, and squirrel made up 75% of the contribution to PC2 (Appendix S1: Figure S6). In 2019, the first two axes accounted for 80.1% of the total variance; cat, raccoon, groundhog contributed 68% to PC1 and squirrel, mouse, and deer contribute 86% towards PC2. In 2018, park connectivity correlated most strongly along the second dimension and park area and Shannon index correlated diagonally between the first and second dimension in opposite directions. In 2019, park area correlated with both PC1 in the negative direction and PC2 in the positive direction and park connectivity showed negative correlation with PC1 and PC2. The Shannon index was correlated diagonally between the first and second dimension and showed positive correlation with both PC1 and PC2 in 2019.

### Tick abundance models

Park area and connectivity had a significant positive effect on DON and there was no significant relationship between Shannon index and DON (Figure 2). The best fit multivariate GLM model that examined the effect of host activity in 2018 on *I. scapularis* density of nymphs in 2019 showed a significant positive effect of deer activity (ß = 1.31, p < 0.001) and mouse abundance (ß = 0.47, p < 0.001), while raccoon activity had a significant negative effect on nymphal abundance the following year (ß = -0.29, p = 0.01, Figure 3.).

**Figure 2.**
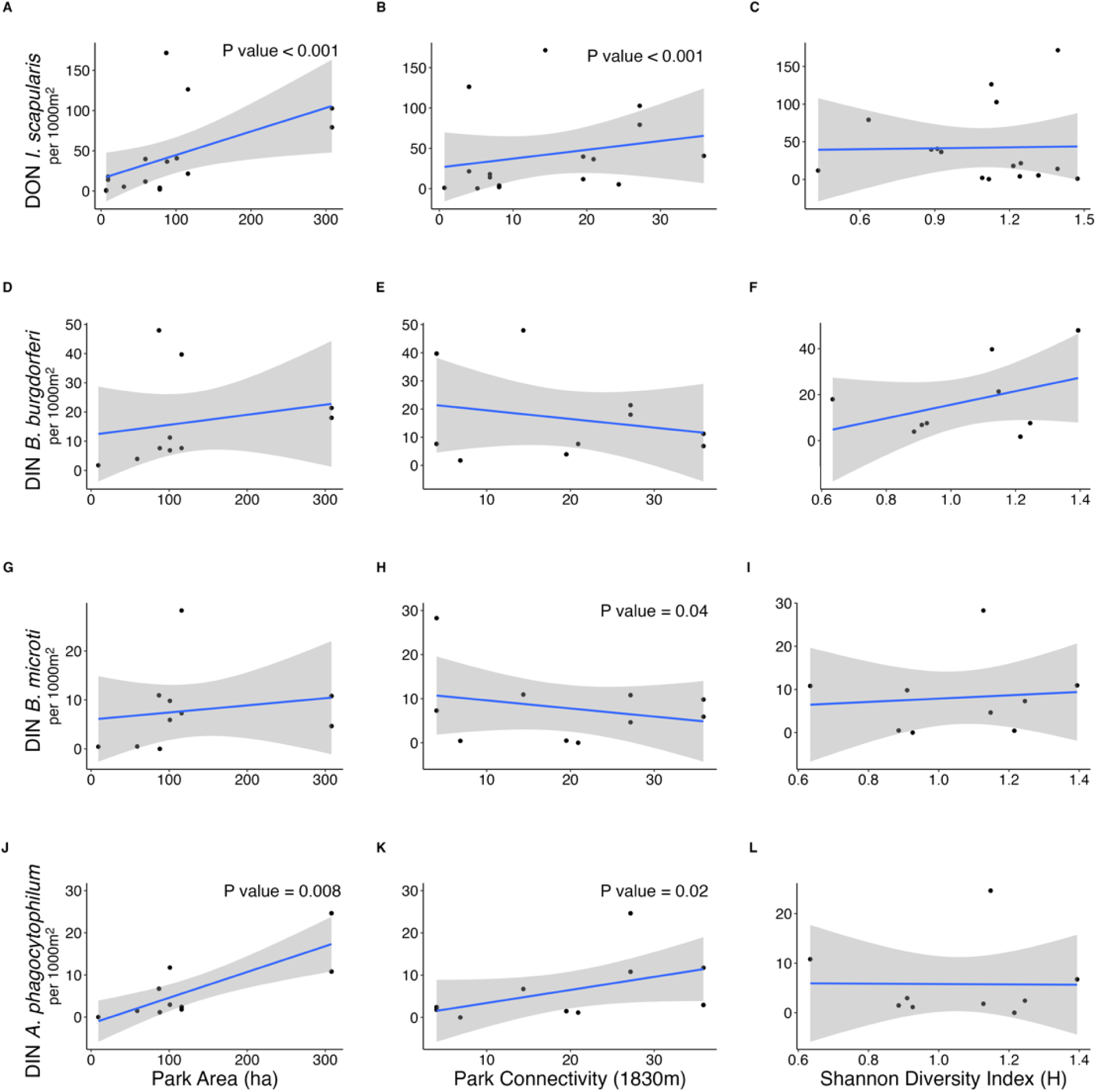
Relationships between park characteristics and the density of *I. scapularis* nymphs (DON) and the density of infected *I. scapularis* nymphs (DIN) with *B. burgdorferi*, *B. microti*, and *A. phagocytophilum.* The P-values listed indicate the significant relationships resulting from negative binomial GLMs. If no P-value is shown in the plot, the relationship was not significant. The blue line shows the smoothed mean with a 95% confidence interval band in gray. The connectivity variable used is the mean cumulative flow within a 1830m moving window.

**Figure 3.**
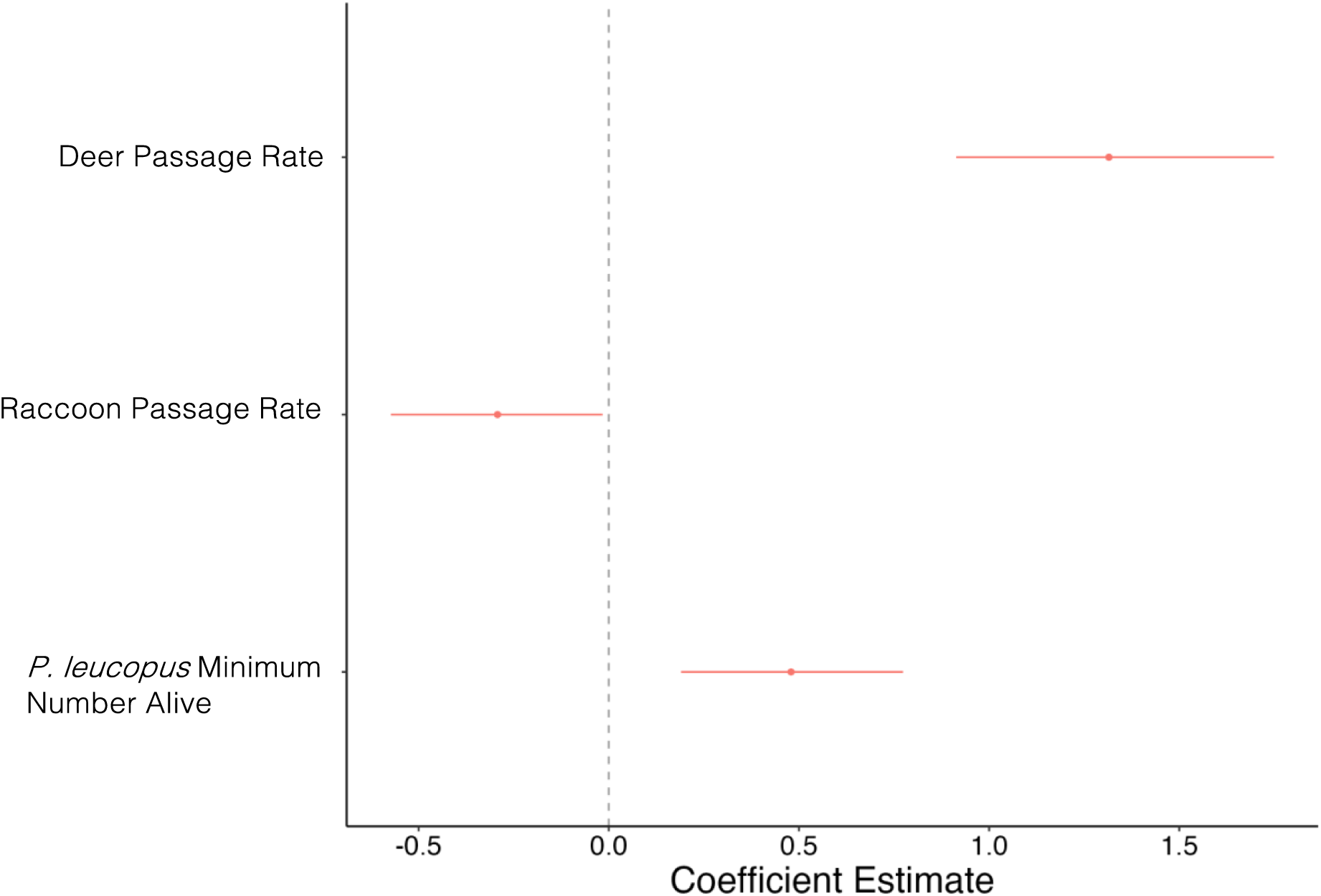
Coefficient plot showing the effect of host passage rate and *P. leucopus* minimum number alive in 2018 on the density of nymphs in 2019.

### Density of infected nymphs models

We did not identify any significant relationships between the density of nymphs infected with *B. burgdorferi* and park characteristics. Park connectivity had a significant negative effect on DIN with *B. microti* while park area and connectivity had significant positive effects on DIN with *A. phagocytophilum* (Figure 2).

### Nymphal infection prevalence models

The best fit model using host encounter probabilities to predict *B. burgdorferi* NIP included deer encounter probability which had a significant negative effect on the NIP (Table 3. and Figure 4A.) and was consistent across all models. We identified significant associations between host encounter probabilities and *B. microti* and *A. phagocytophilum* NIP (Table 3.). The best fit model predicting *A. phagocytophilum* NIP included deer and raccoon encounter probability, where deer showed a significant positive effect on *A. phagocytophilum* NIP and raccoon showed a negative effect on *A. phagocytophilum* NIP (Table 3. and Figure 4B.). The second ranking model predicting NIP with *A. phagocytophilum* showed significant positive effects from the encounter probability of deer and mouse abundance. The best fit model predicting *B. microti* NIP included mouse abundance which had a significant positive effect on *B. microti* NIP estimates (Table 3. and Figure 4C.).

**Figure 4.**
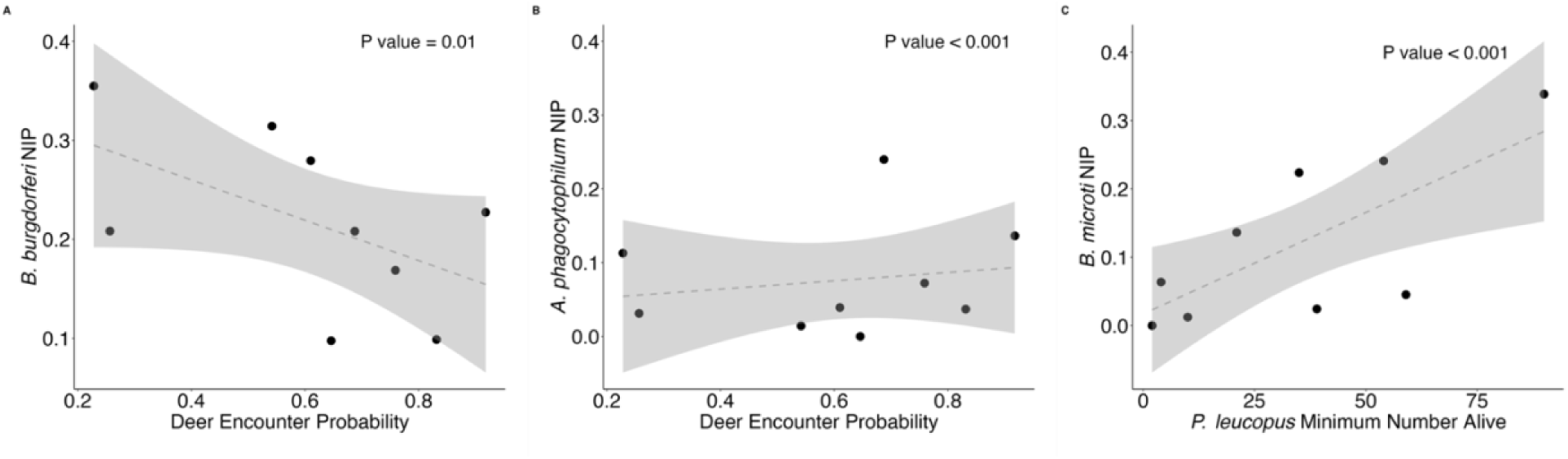
Logistic regression models of nymphal infection prevalence (NIP) of *B. burgdorferi, A. phagocytophilum,* and *B. microti* predicted by host abundance and activity in 2018 and 2019. The best fit models resulted in (A) deer encounter probability as a negative predictor of *B. burgdorferi* NIP and (B) a positive predictor for *A. phagocytophilum* NIP and *P. leucopus* minimum number alive as a positive predictor of *B. microti* NIP (C).

**Table 3.** Model selection table showing the coefficient estimates and the delta AIC for the binomial GLMs of 2018 and 2019 host encounter probabilities (EP) predicting 2018 and 2019 nymphal infection prevalence with *B. burgdorferi, A. phagocytophilum,* and *B. microti*.

| *Borrelia burgdorferi* nymphal infection prevalence |  |  |  |  |
| --- | --- | --- | --- | --- |
| Intercept | Deer EP | Raccoon EP | MNA | ΔAIC |
| **-1.1549\*\*\*** | **-0.2065\*** | - | - | 0 |
| **-1.1632\*\*\*** | **-0.2124\*\*** | -0.0813 | - | 0.49 |
| **-1.1525\*\*\*** | **-0.2031\*** | - | 0.0284 | 1.87 |
| **-1.650\*\*\*** | **-0.2147\*\*** | -0.0865 | -0.0150 | 2.46 |
| *Babesia microti* nymphal infection prevalence |  |  |  |  |
| Intercept | Deer EP | MNA |  | ΔAIC |
| **-3.0585\*\*\*** | - | **2.0146\*\*\*** |  | 0 |
| **-2.6177\*\*\*** | -0.4187 | **1.5523\*\*\*** |  | 0.14 |
| *Anaplasma phagocytophilum* nymphal infection prevalence |  |  |  |  |
| Intercept | Deer EP | Raccoon EP | MNA | ΔAIC |
| **-2.405\*\*\*** | **0.560\*\*** | **-0.260 .** | - | 0 |
| **-2.623\*\*\*** | **0.810\*\*\*** | - | **0.971\*\*\*** | 0.63 |
| **-2.355\*\*\*** | **0.579\*\*\*** | - | - | 1.58 |
| **-2.362\*\*\*** | **0.506\*** | -0.338 | -0.230 | 1.87 |

The top models shown are within 2 delta AIC of the lowest AIC model. Site sample sizes for 2018 and 2019 host data and NIP data = 9. Squirrel EP was not included in the global models due to high multicollinearity values. All remaining covariate VIF scores are under 3. Significant coefficients are shown in bold with the following significance levels: 0 (\*\*\*), 0.001(\*\*), 0.01 (\*), 0.1 (.)

## Discussion

This study used a community ecology approach to understand the determinants and impact of host community assemblage on *I. scapularis* and tick-borne pathogen prevalence in highly fragmented urban green spaces. We first examined how habitat amount and connectivity structured the host community, then assessed the cascading effect of the host community on tick abundance and infection with multiple *I. scapularis*-borne pathogens. Our findings challenge a simple association between forest fragmentation and host community composition. We found that while white-tailed deer are consistently dependent on landscape connectivity, white-footed mice abundance was negatively associated with landscape-scale connectivity and mice were more abundant in larger and more host diverse sites. However, it is important to note annual fluctuations of mice populations in response to resource fluxes are extremely common (Elias et al. 2004, Marcello et al. 2008) and this pattern should be examined over longer periods of time. Secondly, we found that human-adapted species, gray squirrels and raccoons, dominate in the most fragmented settings (smaller, isolated parks) which contrasts an underlying assumption of the dilution effect that decreasing habitat patch size is linked with dominance of competent hosts for *B. burgdorferi* transmission, in particular, white-footed mice.

We observed associations between landscape metrics and the density and infection of nymphs that we hypothesize to be an indirect effect of the host community response to habitat connectivity and composition. We observed lower DON at smaller and more isolated park patches. This contrasts with findings from Allan et al. (2003), where smaller patches had higher DON. However, because our smaller patches were similar in size to ‘large’ patches from Allan et al. (2003) and that study does not report connectivity, comparisons are difficult. Because smaller patches had lower numbers of nymphs, we could not obtain a robust estimate of NIP at some sites, challenging our ability to assess the association between reservoir host community and infection, as found in previous studies (LoGiudice et al. 2008, Diuk-Wasser et al. 2021). More connected and larger parks significantly predicted DIN with *A. phagocytophilum* where larger parks had higher densities of infected nymphs. This may be due to high deer activity in connected parks amplifying the abundance of host seeking nymphs the following year through providing bloodmeals to larva and, because deer are reservoir hosts for *A. phagocytophilum*, leading to higher infection prevalence of nymphs with *A. phagocytophium*. Further, the negative association between DIN with *B. microti* and park connectivity is likely moderated by the lower abundance of mice in highly connected parks, as white-footed mice are the primary host of *B. microti*.

Cascading effects of host activity on infection prevalence were detected for host-limited pathogens and forest-dependent species. The positive association of *B. microti* NIP with white-footed mice and *A. phagocytophilum* NIP with white-tailed deer and white-footed mice, respectively, is consistent with host specialization by *B. microti* and *A. phagocytophilum*, i.e. fewer host species involved in driving their enzootic transmission, as supported by prior studies (Massung et al. 2003a, Yabsley and Shock 2013). Specifically, *B. microti* is associated with a higher abundance of white-footed mice (Spielman et al. 1981, Mather et al. 1990, Goethert et al. 2018). There were positive associations between *A. phagocytophilum* NIP and white-tailed deer and mice activity making it challenging to hypothesize which variant may be circulating since deer are the primary reservoir host of Ap-variant 1 (Massung et al. n.d.) and white-footed mice are reservoir hosts for Ap-ha. Research on the specific genotypes circulating in our study area would yield critical insight into *A. phagocytophilum* host association.

We did not observe significant relationships between landscape characteristics and DIN with *B. burgdorferi* likely due to *B. burgdorferi*’s wider host range. However, the significant negative relationship between deer activity and NIP with *B. burgdorferi* supports the role of deer as a pathogen dilution host for *B. burgdorferi* as has been previously shown (Luttrell et al. 1994, Huang et al. 2019, Pearson et al. 2023, Goethert et al. 2023). We should note that our study design did not account for all possible host species. For example, bird species and chipmunks were rarely captured in photos on our grid sites, and shrews were trapped only at a small subset of sampling sites, leaving us unable to account for the relative role of these host species.

Our study indicates that the drivers of host community composition in fragmented habitat patches are fundamentally different in urban areas in the Northeast U.S. from those observed in agricultural landscapes in the Midwest (Nupp and Swihart 1998, 2000), which were traditionally cited in the literature describing the effects of habitat fragmentation on tick-borne disease risk. While those studies identified a nested community assembly pattern dominated by white-footed mice in small and isolated fragments, our results show dominance by raccoons and squirrels, the most human-dependent hosts, in these patches. This finding indicates that anthropogenic resource subsidies (gardens, trash, supplemental feeding) present in the urban matrix surrounding isolated forest patches influence the host community composition (Rodewald et al. 2011). In fact, when comparing species richness between residential yards and bordering parks on Staten Island, yards had higher mammal species richness and higher detection of low reservoir competent (“dilution”) hosts for *Borrelia burgdorferi*, such as opossums, than paired forested greenspaces (Bastard et al. 2024), potentially driven by resource provisioning.

Although all the species considered here are generally human-adapted, they vary in their degree and type of dependence on the anthropogenic niche (Hulme-Beaman et al. 2016). Due to their body mass and natural history, deer rely on vegetation for forage, safe movement, to provide cover when with young, and during seasonal reproductive activities (VanAcker et al. 2023). Raccoons, however, can persist within the urban matrix without relying on wooded habitat fragments (Gross et al. 2012) and are positively associated with human anthropogenic subsidies (Reed and Bonter 2018), more so than white-footed mice which generally avoid human activity (Hummell et al. 2023). In an across-city comparison of mammalian species occupancy, raccoons and gray squirrels had the highest occupancy across 20 North American cities while white-tailed deer showed a strong negative response to urban intensity (Magle et al. 2021). There was, however, a nonlinear association between urbanization and raccoon and gray squirrel occupancy, with both displaying a positive response to the city’s average housing density only after a threshold of % greenspace was reached (squirrels) and before a maximum housing density was reached (raccoons) (Fidino et al. 2021). Future work should investigate how shifts in the host community composition at these thresholds across cities cascade to impact tick-borne disease risk. In the PCA, we observed the ordination of raccoon along the two axes change directions between the years indicating that this species may exhibit higher plasticity in its response to anthropogenic resource subsides, which are temporally and spatially variable, or that small sample sizes led to inconsistent results. Thus, although all the species studied can be found in an urban landscape, some species are more tightly limited by ecological features such as vegetation and forest cover or to the total habitat amount on the landscape, while others rely more strongly on anthropogenic food resources.

We determined higher rates of nymphal coinfection with *B. burgdorferi* and *B. microti* than expected by chance, supporting previous work that identified this coinfection pattern in field-derived *I. scapularis* nymphs (Prusinski et al. 2014, Diuk-Wasser et al. 2016). Since high proportions of *B. burgdorferi* infection in *P. leucopus* populations lowers the ecological threshold for *B. microti* invasion (Dunn et al. 2014), we can expect sites with prior establishment of *B. burgdorferi* to experience earlier invasion with *B. microti*. The discovery of vertical transmission of *B. microti* in *P. leucopus* also indicates that sites with higher *P. leucopus* populations would enhance *B. microti* pathogen pressure to feeding ticks (Tufts and Diuk-Wasser 2018). To reduce the hazard of *I. scapularis* coinfected with *B. burgdorferi* and *B. microti* for humans, vaccinating *P. leucopus* populations for *B. burgdorferi* may reduce the transmission of *B. microti* alongside the primary target of *B. burgdorferi* (Vannier et al. 2023). Finally, reducing populations of *P. leucopus* could reduce the prevalence of *B. microti* in mice through interrupting horizontal and vertical transmission events.

One of the largest barriers to quantifying pathogen transmission within a community framework is the need to trap and sample live hosts to measure the host’s ability to feed ticks and transmit pathogens. However, live trapping rarely provides large enough samples to assess host population sizes and does not provide information on host movement. We have shown how remote monitoring through camera trapping can be used to estimate a metric of host availability to questing ticks that integrates measures of host abundance and activity. Combined with host trapping to assess hosts’ tick burdens and infection along with tick sampling (Haydon et al. 2002, Viana et al. 2014, Takumi et al. 2019), this method can efficiently detect patterns of host community composition as its directly relevant to questing ticks within a specific ecological context of interest.

*Ixodes scapularis* continues to expand throughout North America and now occupies new urban expansion fronts allowing for studies such as ours to empirically assess the previously modeled links between landscape, host community composition, and tick abundance and tick-borne pathogen prevalence. We have demonstrated that with extreme habitat fragmentation, anthropogenic resource subsidies and species-specific responses to anthropogenic pressures shifts the host community to become dominated by human-adapted hosts (Fig. 5. Fidino et al. 2021), typically considered dilution hosts for *B. burgdorferi* transmission, such as raccoons and squirrels rather than forest-dependent hosts like deer and white-footed mice. This change in host community alters the relative abundance of bloodmeals available to questing ticks which may relate to the local extinctions of tick and pathogen populations we observed in highly isolated habitat patches. At extreme levels of fragmentation, tick-borne pathogens become extinct without an abundance of permissive hosts for bloodmeals to support tick populations for ongoing pathogen transmission. Our work documents this process with pathogens such as *B. microti* and *A. phagocytophilum*, which are more vulnerable to extinction with high fragmentation due to the pathogens’ narrow host range and lower transmission efficiency.

## Acknowledgements

The authors would like to especially thank Katherine Uriarte, Mandy Weaver, Nicole Pietrunti, Laura Plimpton, and Ted Chritton for help with data collection. This publication was supported by the Cooperative Agreement Number U01CK000509-01 between the Centers for Disease Control and Prevention and Northeast Regional Center for Excellence in Vector Borne Diseases. Its contents are solely the responsibility of the authors and do not necessarily represent the official views of the Centers for Disease Control and Prevention, the Department of Health and Human Services or the National Science Foundation. Funding was also provided by the National Science Foundation’s Coupled Natural Human Systems 2/Dynamics of Integrated Socio-Environmental Systems (CNH2/DISES) program (Award #1924061).

