## Supplemental File for "Urban greenspace connectivity drive shifts in host assemblages and tick-borne pathogen infection"

**Appendix S1**

Shifts in urban greenspace connectivity drive host assemblages and tick-borne pathogen prevalence

**Authors:**
Meredith C. VanAcker[1,2], Tim R. Hofmeester[3], Jeffrey Zhang-Sun[1], Heidi K. Goethert[4], Maria A. Diuk-Wasser[1]

**Section S1: Methods**

DNA Extraction of *Ixodes scapularis* nymphs
Each nymphal tick was placed at the bottom of a 1.7 ml microcentrifuge tube and dried if the tick was preserved in ethanol. The tube was placed in liquid nitrogen for 5 seconds then the tick was pulverized using a plastic pestle. If nymphs were frozen at -80°C, the tubes were directly placed in liquid nitrogen and pulverized. All buffers and reagents were sourced from Qiagen DNeasy Blood and Tissue kit (Qiagen, Valencia, CA, USA). 180 µL ATL buffer and 25 µL of Proteinase K were added to each tube, vortexed for 15 seconds and centrifuged for 30 seconds. All tubes were incubated overnight at 56°C. Each sample was briefly centrifuged and then loaded into the Qiagen QIAcube HT DNA extraction system (Qiagen).

DNA Extraction of *Peromyscus leucopus* blood
Three to four 3mm punches of dried blood blot were punched from the Whatman FTA cards and placed into a 1.7 ml microcentrifuge tube. 280 µL ATL buffer and 20 µL of Proteinase K were added to each tube and vortexed for 5 seconds (Qiagen, Valencia, CA, USA). The samples were incubated at 56°C for 2 hours, mixed every 30 minutes, and centrifuged following incubation. 300 µL AL buffer was added, the sample was vortexed for 10 seconds, and incubated at 70°C for 10 minutes, mixing every 3 minutes. The samples were centrifuged and then loaded into the Qiagen QIAcube HT DNA extraction system (Qiagen).

Pathogen Screening
All *I. scapularis* ticks collected from each site were screened individually for infection with *B. burgdorferi*, *B. microti*, and *A. phagocytophilum* using a multiplex real-time PCR assay with primers and probes that were previously published(Tokarz et al. 2017). The plasmid-derived gene *ospA* was used as a target for *B. burgdorferi*, the mitochondrial *cox-1* gene and the 16S rRNA gene were used to target *B. microti* and *A. phagocytophilum*, respectively (Tokarz et al. 2017). DNA samples were amplified in 15 µL reaction mixtures which included 1.5 µL tick DNA and 13.5 µL Multiplex Powermix (Bio-Rad Laboratories, Hercules, CA) using a Light Cycler 480 real-time PCR machine (Roche, Basel, Switzerland). The PCR reaction conditions were 98°C for 30 seconds and then 50 cycles of 98°C for 8 seconds and 58°C at 30 seconds using the primers listed in Table S2.2. Single samples were run and considered positive if amplification peaked by cycle 44. If no amplification signal was detected the sample was determined to be negative. PCR controls were included in every run to detect potential cross-contamination.

Each *P. leucopus* blood sample was tested in duplicate for the presence of *B. microti*. Samples were run on a 7500 real-time PCR system (Applied Biosystems®, ThermoFisher

Scientific, Waltham, WA, USA) using TaqMan Fast Advanced chemistry (ThermoFisher Scientific, Waltham, WA) and cycling conditions consisted of: 95 °C for 20 seconds, followed by 40 cycles of 95 °C for 3 seconds and 60 °C for 30 seconds using the primers listed in Table S2.3 (Tufts and Diuk-Wasser 2018). The average cycle threshold and quantity values were retrieved from each run and used to determine the infection status of each sample. PCR controls were included in every run to detect potential cross-contamination.

Camera deployment

The camera settings were configured for high PIR sensitivity (high motion sensitivity), 3 picture burst, and used a 30 second detection delay to reduce capturing repetitive photos of few individuals and to preserve battery life and memory card storage space. Calibration photos were set to be automatically taken at 00:00 and 00:30 daily, which allowed for a camera functionality check by assessing photo quality during photo downloads. Every two weeks and within 24 hours of the beginning of each sampling session, we downloaded photos, inspected cameras for functionality, and rotated the cameras' location within the grids to a new random set of coordinates.

Connectivity Layer

We calculated a metric called cumulative current by using the program Omniscape (Landau et al. 2021) to assess the importance of deer selected habitat types in maintaining connectivity across SI for deer. We used previously estimated beta coefficients resulting from individual step selection analysis conducted from Staten Island deer GPS data (VanAcker et al. 2023) to inform the raster pixel resistance values. Thus, we assigned resistance values to each land cover class on Staten Island according to deer's relative selection strength for a particular land cover class in reference to forest land cover. Specifically, we averaged the beta coefficients for each land cover class weighted by the coefficient estimate's inverse variance. These weighted averages were rescaled to range from 1-1000 and assigned to the six land cover classes to be used as resistance values. We applied Omniscape which runs Circuitscape iteratively through the landscape in a moving window within a user assigned radius to evaluate connectivity between every possible pair of pixels that are no farther apart than the moving window distance (Landau et al. 2021). We ran Omniscape with four different moving window radii that reflected Staten Island deer movement where 120m was roughly equivalent to the distance that deer responded by high intensity movement, 480m, 990m, and 1830m was roughly equivalent to the largest home range area for male deer (VanAcker et al. 2023). The block value was standardized to 10% of the radius value provided (Belote et al. 2022).

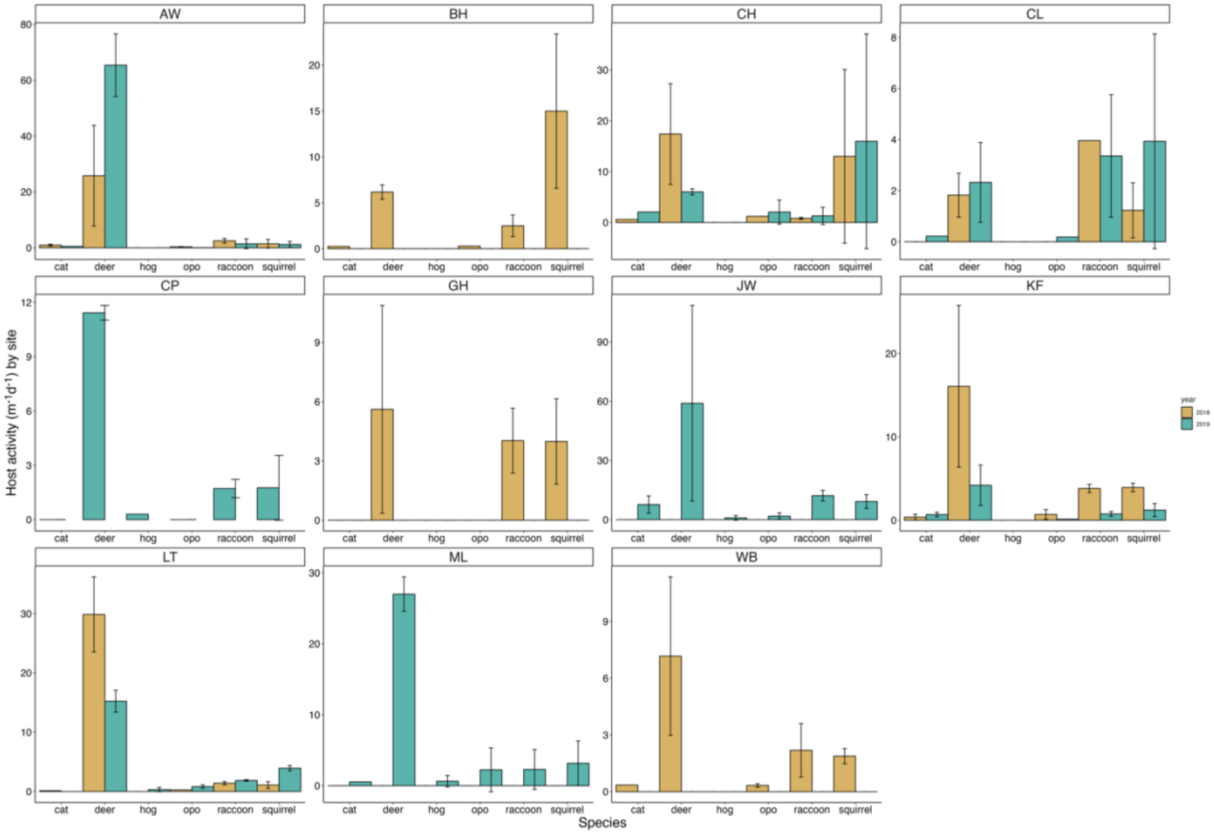

**Figure S1.** The estimated passage rate (PR) per site for each species in 2018 and 2019. The host species name is along the x-axis where opo is an abbreviation for opossum and hog is an abbreviation for groundhog. The error bars indicate the standard deviation of the passage rate estimates across the two cameras present per site/year. Gold bars indicate 2018 PR and teal bars represent 2019 PR. Note the different scaling of the y-axes and that not all species were detected at each site. Five sites were sampled both years.

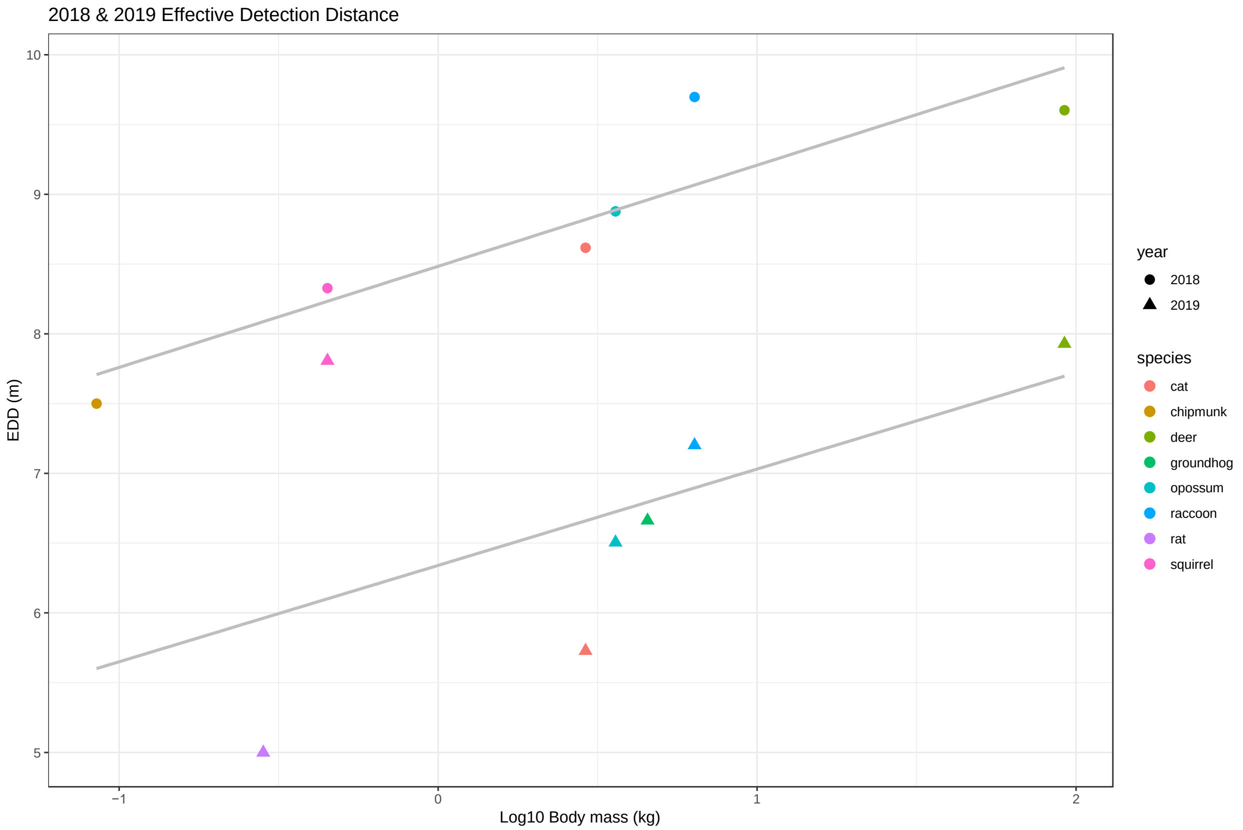

**Figure S2.** The relationship between estimated effective detection distances (EDD) for each host species and body size by camera trap year.

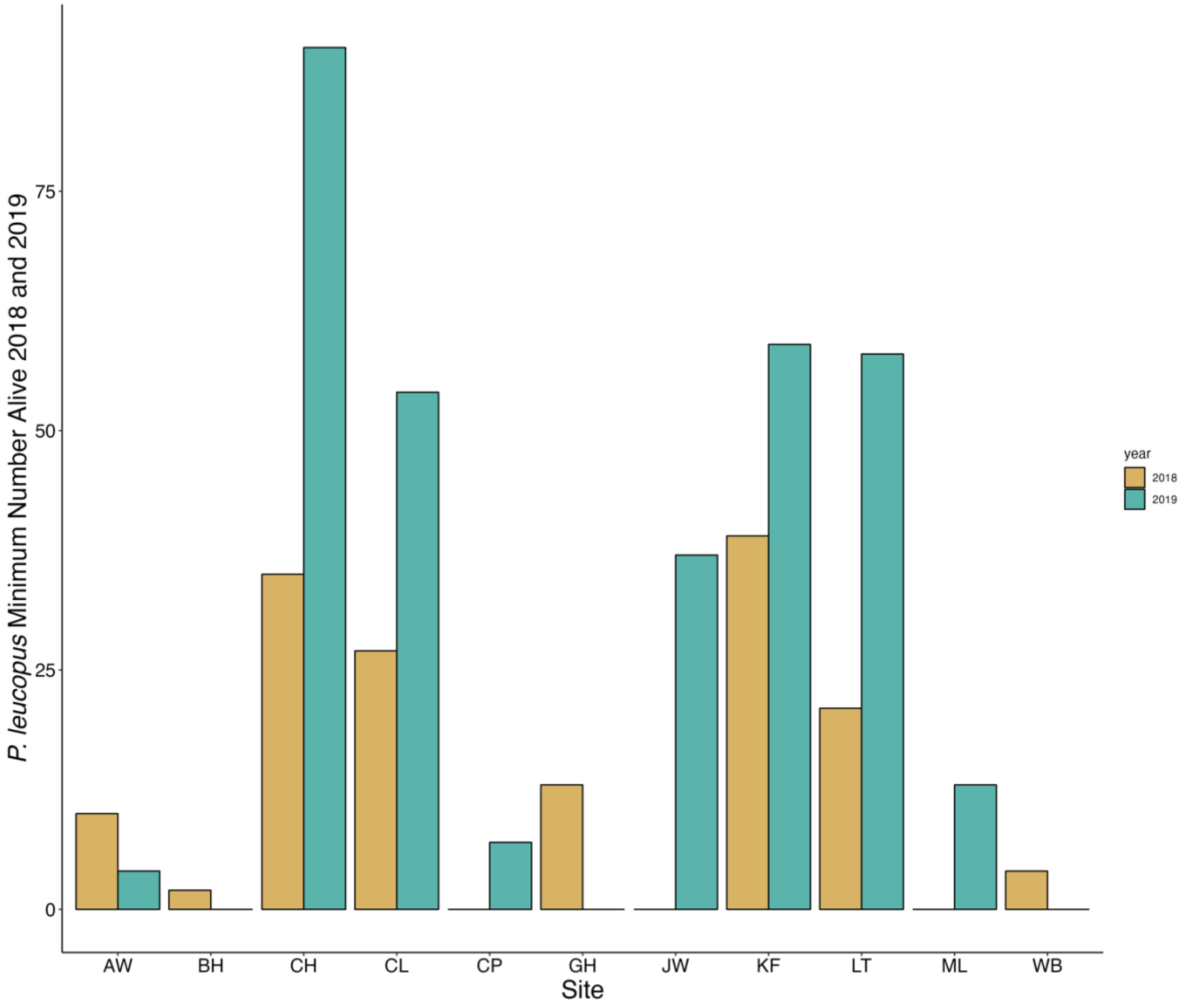

**Figure S3.** The minimum number alive for *P. leucopus* per site in 2018 and 2019.

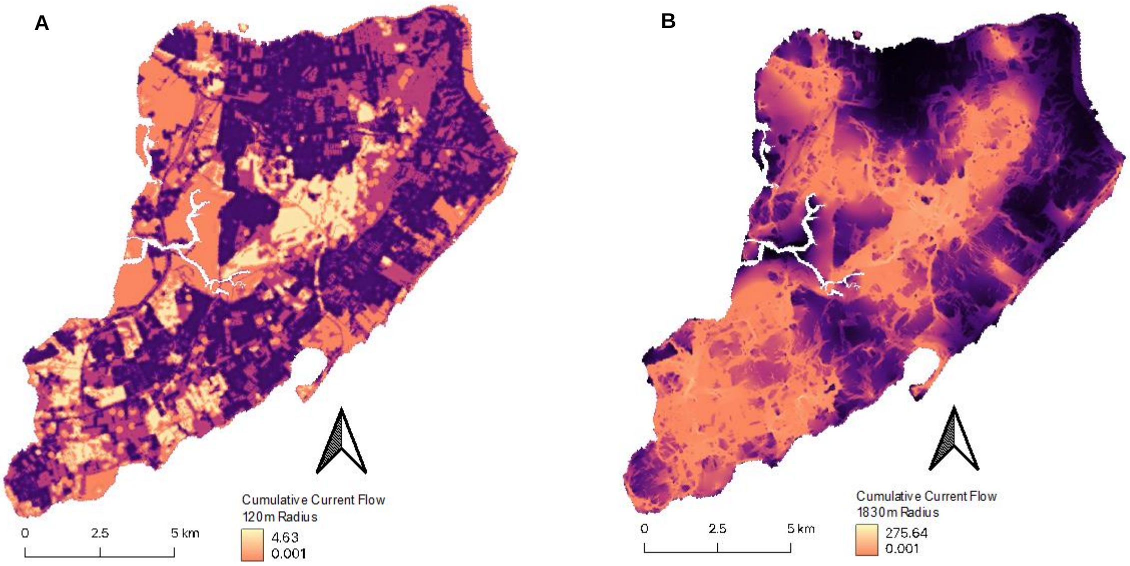

**Figure S4.** Maps displaying the best fit moving window scales from Omniscape for predicting deer passage rate (A) and the density of nymphs (DON) (B). The median cumulative current within 120m was the strongest predictor of deer passage rate while the mean cumulative current flow with 1830m moving window was the best predictor of DON.

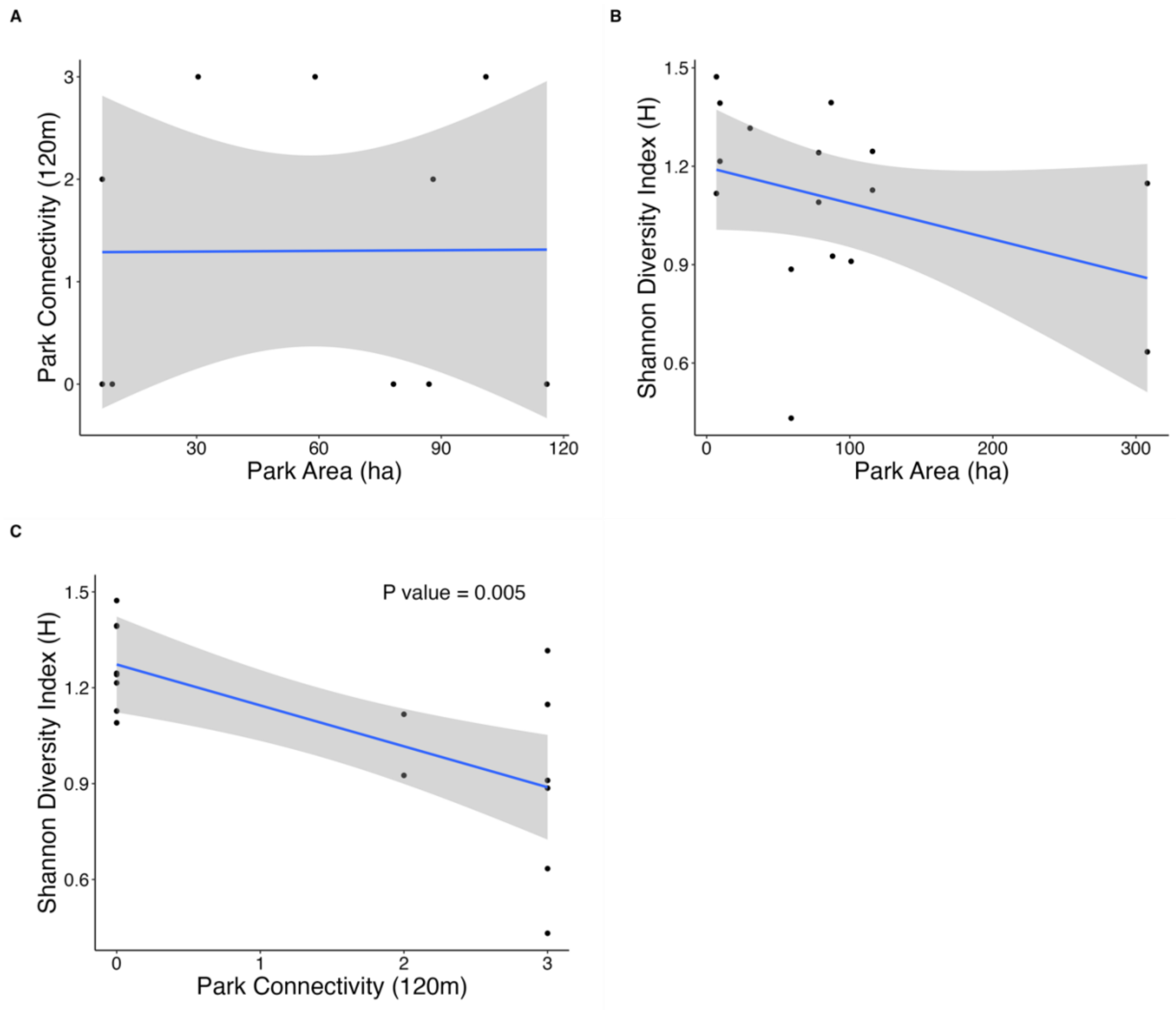

**Figure S5.** Univariate relationships between park area, connectivity, and Shannon's diversity index. Park area and connectivity values remain static between the two years and Shannon's diversity index varies between years. Blue lines show the model fit of each univariate gaussian model using the GLM smoothing method and the gray bands indicate 95% confidence intervals. The P-value is included for the significant model.

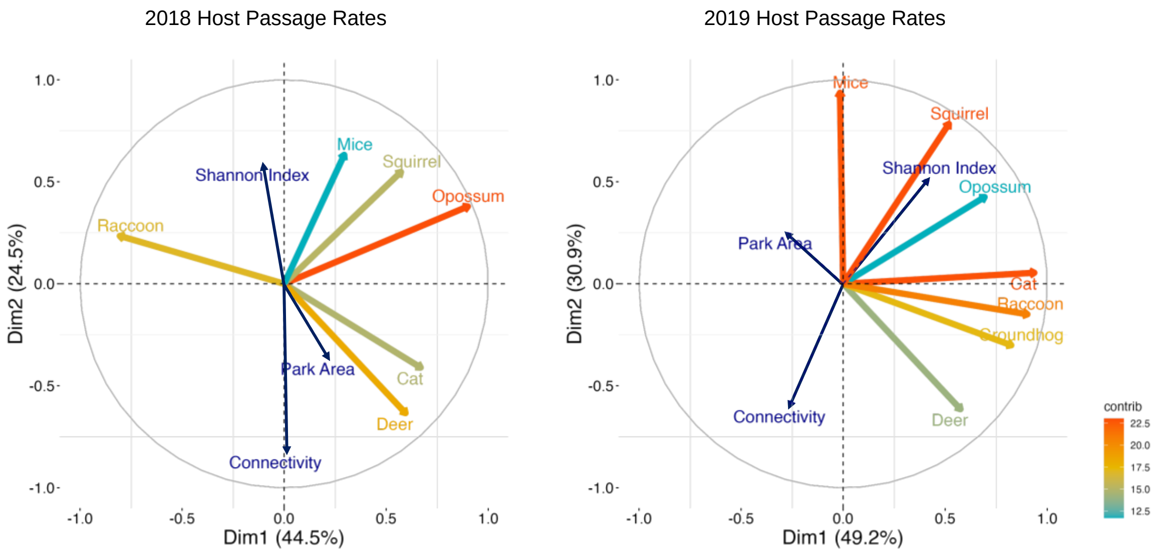

**Figure S6.** Principal components analysis of host passage rates and park characteristics in 2018 and 2019. The x-axis shows principal component 1 and the y-axis shows principal component 2 with the percent variation in the data explained by each axis in parentheses. The color of each line indicates a host passage rate contribution to principal axes. The dark blue lines show the park-level covariates where the lines' location over coordinate space is determined by the correlation between the respective park characteristic and the principal component axes and the line's length shows how well the variable is represented by the two components. If the variable is closer to circle of correlations, the variable can be better reconstructed from the first two components and if the variable is closer to the center of the plot, it is less important for the first two components. The connectivity variable used is the median cumulative flow within a 120m moving window.

**Table S1. (a)** Primer and probe sequences used for *I. scapularis* pathogen screening (Tokarz et al. 2017)

| **Agent** | **Gene target** | **Forward primer** | **Reverse primer** | **Probe**[a] |
|---|---|---|---|---|
| *Borrelia burgdorferi* | ospA | CCTTCAAGTACTCCAGATCCATTG | AACAAAGACGGCAAGTACGATC | FAM-CAACAGTAGACAAGCTTGA-MGB |
| *Babesia microti* | cox1 | CATCATGCCAGGCCTGTTTG | GAAGAAACCACAAGAGCAAATGC | Quasar 705-TACTACCCATACTGGTCGGTGCTCC-BHQ |
| *Anaplasma phagocytophilum* | 16S rRNA | GGCATGTAGGCGGTTCGGT | CACTAGGAATTCCGCTATCCTCTCC | Cy5-GCCAGGGCTTAACCCTGGAGCT-BHQ |

[a]Abbreviations: FAM, 6-carboxyfluorescein; MGB, minor groove binder; BHQ, black hole quencher dye.

**(b)** Primer and probe sequences used for *P. leucopus* pathogen screening (Rollend et al. 2013)

| **Agent** | **Gene target** | **Forward primer** | **Reverse primer** | **Probe** |
|---|---|---|---|---|
| *Babesia microti* | 18S rRNA | AACAGGCATTCGCCTTGAAT | CCAACTGCTCCTATTAACCATTACTCT | 6FAM-CTACAGCATGGAATAATGA-MGBNFQ |

| Year | Site | No. PCR + (% Infection Prevalence) | | | | Coinfection | | |
|---|---|---|---|---|---|---|---|---|
| | | *B. burgdorferi* | *B. microti* | *A. phagocytophilum* | Total no. tested | Bb/Bm | Bb/Ap | Bm/Ap |
| 2018 | Arden Woods* | 8 (9.88%) | 1 (1.23%) | 3 (3.70%) | 81 | 0 | 0 | 0 |
| 2019 | Arden Woods | 2 (25%) | 0 | 0 | 8 | 0 | 0 | 0 |
| 2018 | Blue Heron* | 20 (20.83%) | 0 | 3 (3.13%) | 96 | 0 | 0 | 0 |
| 2019 | Blue Heron | 3 (12%) | 1 (4%) | 0 | 25 | 0 | 0 | 0 |
| 2018 | Clay Pit* | 25 (12.5%) | 14 (7%) | 26 (13%) | 200 | 4 | 2 | 1 |
| 2019 | Clay Pit* | 14 (16.87%) | 20 (24.10%) | 6 (7.23%) | 83 | 5 | 0 | 1 |
| 2018 | Clove Lakes | 0 | 0 | 0 | 4 | 0 | 0 | 0 |
| 2019 | Clove Lakes | 0 | 0 | 0 | 10 | 0 | 0 | 0 |
| 2018 | Conference House* | 66 (31.43%) | 47 (22.38%) | 3 (1.43%) | 210 | 31 | 1 | 0 |
| 2019 | Conference House* | 22 (35.48%) | 21 (33.87%) | 7 (11.29%) | 62 | 12 | 4 | 2 |
| 2018 | Goodhue | 0 | 0 | 0 | 1 | 0 | 0 | 0 |
| 2018 | King Fisher* | 4 (9.76%) | 1 (2.44%) | 0 | 41 | 0 | 0 | 0 |
| 2019 | King Fisher | 1 (20%) | 0 | 0 | 5 | 0 | 0 | 0 |
| 2018 | Latourette* | 45 (22.73%) | 27 (13.64%) | 27 (13.64%) | 198 | 14 | 4 | 2 |
| 2019 | Latourette* | 46 (20.81%) | 10 (4.52%) | 53 (23.98%) | 221 | 2 | 2 | 3 |
| 2019 | Mount Loretto | 0 | 0 | 0 | 4 | 0 | 0 | 0 |
| 2018 | Willowbrook* | 57 (27.94%) | 13 (6.37%) | 8 (3.92%) | 204 | 7 | 0 | 1 |
| 2019 | Willowbrook | 5 (41.67%) | 0 | 1 (8.33%) | 12 | 0 | 0 | 0 |
| 2018 | Overall | 225 (21.31%) | 103 (9.75%) | 70 (6.63%) | 1056 | 56 | 7 | 4 |
| 2019 | Overall | 105 (21.88%) | 55 (11.46%) | 67 (13.96%) | 480 | 19 | 6 | 6 |

**Table S2.** *I. scapularis* nymphal infection prevalence and coinfection for *B. burgdorferi*, *B. microti*, and *A. phagocytophilum* in 2018 and 2019 from all sites sampled. In 2018 there were 56 *I. scapularis* nymphs that were coinfected with *B. burgdorferi* and *B. microti*, 7 nymphs coinfected with *B. burgdorferi* and *A. phagocytophilum*, and 4 nymphs coinfected with *B. microti* and *A. phagocytophilum* out of 1056 nymphs screened. In 2019 there were 19 nymphs that were coinfected with *B. burgdorferi* and *B. microti*, 6 nymphs coinfected with *B. burgdorferi* and *A. phagocytophilum*, and 6 nymphs coinfected with *B. microti* and *A. phagocytophilum* out of 480 nymphs screened. Each year one nymph tested positive with all three pathogens. Nymphs coinfected with *B. burgdorferi* and *B.*

*microti* occurred more often that what would be expected by chance (2018: $X^2 = 55.81$; $P < 0.0001$; 2019: $X^2 = 49.146$; $P < 0.0006$) across both years. Coinfection with either *B. burgdorferi* or *B. microti* and *A. phagocytophilum* was not different than predicted by chance. The '*' indicates the site-years included in all GLM modeling as the total number of ticks tested is greater than 40.

| Response variable | Intercept | Park connectivity | Park area | Shannon Diversity | ΔAIC |
|---|---|---|---|---|---|
| Deer PR | **-1.4911*** ** | **0.1263*** | **-0.2930*** ** | **-0.2684*** ** | 0 |
| Raccoon PR | **-3.3105*** ** | - | **-0.3763** ** | **0.5295*** ** | 0 |
| Raccoon PR | **-1.7598*** ** | -0.0841 | **-0.5011*** | **0.5553*** ** | 1.8276 |
| Squirrel PR | **-2.6405*** ** | **-0.4760*** ** | **0.1859*** | - | 0 |
| Squirrel PR | **-2.6497*** ** | **-0.4052*** ** | **0.2050*** | 0.1247 | 0.6684 |

**Table S3.** The best fit Poisson Generalized Linear Mixed Models predicting the most dominant host passage rates. The GLMMs listed include any model that falls within 2 ΔAIC from the lowest AIC model. The response variable is the passage rates of deer, raccoon, and squirrels. Park connectivity, area and Shannon diversity are included as predictor models as well as the log of the total number of nights the camera was active as an offset and a random effect for year. PR indicates 'Passage Rate' and (-) means the covariate was not included in the model. The connectivity variable used is the median cumulative flow within a 120m moving window. Significant coefficients are shown in bold with the following significance levels: 0 (***), 0.001(**), 0.01 (*).

| Response variable | Intercept | Park connectivity | Park area | Shannon Diversity | ΔAIC |
|---|---|---|---|---|---|
| MNA | **3.1219*** ** | **-0.7032*** ** | **0.4932** ** | - | 0 |
| MNA | **3.1263*** ** | **-0.6312** ** | **0.4865** ** | 0.0955 | 1.7711 |

**Table S4.** The best fit negative binomial Generalized Linear Mixed Model predicting the minimum number alive (MNA) for *Peromyscus leucopus* for 2018 and 2019. MNA signifies 'minimum number alive' and a random effect for year was used. The connectivity variable used is the median cumulative flow within a 120m moving window. Significant coefficients are shown in bold with the following significance levels: 0 (***), 0.001(**), 0.01 (*).

| Landcover class | Landcover description | Averaged beta coefficient | Rescaled resistance value |
|---|---|---|---|
| Tree canopy | Deciduous, evergreen, and mixed forest | NA | 1 |
| Wetland/Herbaceous/Water | Grassland, herbaceous, woody wetlands, emergent herbaceous wetlands, and barren land | -0.1768 | 97 |
| Open and low intensity development | Open space and low intensity development | -0.3178 | 175 |
| High vegetation residential blocks | Residential development with high vegetation height, low impervious cover, high tree canopy, and large yard areas | -0.3484 | 192 |
| Medium and high intensity development | Non-residential medium and high intensity development | -1.7231 | 950 |
| Low vegetation residential blocks | Residential development with low vegetation height, high impervious cover, low tree canopy, and small yard areas | -1.8134 | 1000 |

**Table S5.** Staten Island landcover class description and resistance values used to produce the Omniscape resistance layer. All landcover classes came from NLCD(U.S. Geological Survey 2019) except for the residential block types which are a composite layer constructed from prior work (VanAcker et al. 2023). The averaged beta coefficient is informed by integrated step selection models using movement data from 27 Staten Island deer between 2016-2021. The beta coefficients were rescaled such that high values indicate highest resistance to current flow and low values indicate the least resistance to current flow.

| | | **Density of nymphs** | | **Deer Passage Rate** | | **Raccoon Passage Rates** | | **Squirrel Passage Rates** | | **MNA** | |
|---|---|---|---|---|---|---|---|---|---|---|---|
| Method of summarization | Omniscape moving window radius (m) | AIC | Delta AIC | AIC | Delta AIC | AIC | Delta AIC | AIC | Delta AIC | AIC | Delta AIC |
| Mean cumulative current | 120 | 3351.08 | 46.44 | 721.42 | 49.6 | 117.96 | **0.0** | 184.00 | **0.0** | 142.81 | 1.8 |
| Mean cumulative current | 480 | 3343.22 | 38.58 | 718.31 | 46.5 | 124.41 | 6.4 | 198.83 | 14.8 | 141.01 | **0.0** |
| Mean cumulative current | 990 | 3329.52 | 24.88 | 719.96 | 48.1 | 128.50 | 10.5 | 206.21 | 22.2 | 141.06 | **0.0** |
| Mean cumulative current | 1830 | 3304.64 | **0.0** | 720.80 | 49.0 | 129.31 | 11.3 | 201.44 | 17.4 | 142.78 | 1.8 |
| Median cumulative current | 120 | 3345.86 | 41.22 | 671.82 | **0.0** | 144.17 | 17.4 | 209.62 | 25.6 | 142.74 | 1.7 |
| Median cumulative current | 480 | 3350.08 | 45.44 | 689.12 | 17.3 | 138.02 | 20.1 | 203.95 | 19.9 | 143.63 | 2.6 |
| Median cumulative current | 990 | 3341.04 | 36.4 | 707.82 | 36.0 | 138.26 | 20.3 | 209.21 | 25.2 | 143.50 | 2.5 |
| Median cumulative current | 1830 | 3330.98 | 26.34 | 718.17 | 46.3 | 135.34 | 26.2 | 203.86 | 19.9 | 143.77 | 2.8 |

**Table S6.** The scale assessment of Omniscape moving window radius ranging from 120m-1830m. The AIC comparison results are from the following negative binomial model structure: DON 2018 & DON 2019 ~ radius scale + (1|year) + offset(log(transect distance)).

| *B. microti* infection prevalence in *P. leucopus* | | | |
|---|---|---|---|
| Year | Sites | No. PCR + (% Infection Prevalence) | Total No. Tested |
| 2018 | Arden Woods | 0 (0%) | 10 |
| 2019 | Arden Woods | 1 (25%) | 4 |
| 2018 | Blue Heron | 0 (0%) | 2 |
| 2019 | Clay Pit | 7 (87.5%) | 8 |
| 2018 | Clove Lakes | 0 (0%) | 26 |
| 2019 | Clove Lakes | 0 (0%) | 40 |
| 2018 | Conference House | 30 (90.9%) | 33 |
| 2019 | Conference House | 50 (74.6%) | 67 |
| 2018 | Goodhue | 1 (9.0%) | 11 |
| 2019 | Jones Woods | 0 (0%) | 26 |
| 2018 | King Fisher | 0 (0%) | 31 |
| 2019 | King Fisher | 11 (29.7%) | 37 |
| 2018 | Latourette | 10 (90.9%) | 11 |
| 2019 | Latourette | 31(79.4%) | 39 |
| 2019 | Mount Loretto | 8 (66.6%) | 12 |
| 2018 | Willowbrook | 0 (0%) | 1 |
| 2018 | Overall | 41 (32.8%) | 125 |
| 2019 | Overall | 108 (45.9%) | 235 |

**Table S7.** *B. microti* infection prevalence for *P. leucopus* from 2018 and 2019.
